# TLR7 Signaling Regulates Embryonic Hematopoietic Stem Cell Development by Sensing microRNA-146a in Vertebrates

**DOI:** 10.64898/2026.07.01.735198

**Authors:** Hongyan Liu, Kun Zhou, Kai Zhu, Yun-Fei Li, Liyuan Mo, Peng-Fei Xu, Yan Li

## Abstract

The specification of hematopoietic stem cells (HSCs) is tightly regulated by multiple transcription factors and signaling pathways. Inflammatory signaling is pivotal for embryonic HSC development, but the mechanisms that activate it *in vivo* remain poorly understood. Here, we show that Toll-like receptor 7 (TLR7) is essential for the emergence of embryonic HSC in both zebrafish and mouse embryos. TLR7 deficiency reduces HSC numbers but not primitive or definitive progenitors. Conversely, the TLR7 agonist R848 enhances embryonic HSC development. Mechanistically, TLR7 signaling acts through interferon regulatory factor 5 (IRF5) to induce the expression of inflammatory cytokines, which subsequently activate Notch signaling to promote HSC emergence through a non-cell-autonomous mechanism. Notably, we identify microRNA-146a (miR-146a) as a potential endogenous activator of TLR7, inducing inflammatory signaling and promoting HSC development. Pharmacological treatment with miR-146a significantly increases HSC numbers in zebrafish embryos. Together, our findings reveal a crucial role for miR-146a-TLR7-IRF5 signaling axis in HSC emergence, providing insights into the endogenous factors that drive tonic inflammatory signaling during normal hematopoiesis and suggesting the translational potential of TLR7 agonists and miR-146a for stem-cell-based therapeutics.

**Significance Statement:** The embryonic origin of hematopoietic stem cells (HSCs) requires inflammatory signals, but the endogenous factor that triggers this process remains elusive. We identify microRNA-146a (miR-146a) as a natural activator of Toll-like Receptor 7 (TLR7) signaling, which is essential for HSC emergence. This miR-146a-TLR7 axis functions through IRF5 and inflammatory cytokines to activate the Notch signaling, specifically promoting embryonic HSC development. Our work addresses the critical question of endogenous ligands that mediate tonic inflammatory signaling in normal hematopoiesis, uncovers novel crosstalk between miRNAs and innate immunity in HSC specification, and identifies promising candidates for stem cell-based therapeutics.

## Introduction

Hematopoietic stem cells (HSCs) are essential for developing and maintaining the hematopoietic system owing to their dual capacity for self-renewal and differentiation into all blood lineages, making them vital for cell-based therapies targeting malignant blood disorders. During embryonic hematopoiesis, HSCs arise from a specialized subset of endothelial cells termed hemogenic endothelial cells (HECs) within the dorsal aorta through endothelial-to-hematopoietic transition (EHT) (1–4). This cellular reprogramming event is orchestrated by diverse intrinsic and extrinsic factors, including transcriptional networks (e.g., RUNX1 and GATA2 (5, 6)), signaling pathways (e.g., Notch and Wnt signaling (7–9)), and mechanical forces (e.g., wall shear stress and cyclic stretch (10, 11)). Among these, inflammatory signaling has been established as a crucial regulator of HSC specification. Multiple pro-inflammatory cytokines, including TNF-α, IFN-α, IFN-γ, IL-1β, IL-6, G-CSF, have been implicated in this process (12–18), while IL-3 is essential for HSC proliferation and maturation (19). Furthermore, pattern recognition receptors such as Toll-like receptor 4 (TLR4), RIG-I-like receptors (RLR), and (NOD)-like receptors (NLRs) have been identified as contributors to HSC emergence (20–22).

A prevailing consensus suggests that the inflammatory signals promoting HSC emergence are from “tonic” inflammation, a state of low-grade, non-pathogenic immune activation, rather than from infection (23). Primitive myeloid cells, including neutrophils and macrophages, are considered important sources of tonic inflammation, as evidenced by neutrophil and macrophage depletion, which decreases the number of hematopoietic stem and progenitor cells (HSPCs) in zebrafish (12, 13). In mice, depleting CD206^+^ macrophages impairs embryonic HSC development (24). Additionally, excessive R-loop accumulation in *Ddx4* mutants enhances HSPC production via cGAS-STING inflammatory signaling (25). Identifying the sources and activators of this tonic inflammation has remained an interesting area of study.

Toll-like receptor 7 (TLR7) is an endosomal RNA sensor expressed in both immune and non-immune cells (26, 27). Upon binding to single-stranded RNA, TLR7 activates transcription factors interferon regulatory factor (IRF) 7 or IRF5, inducing the production of type I interferons (IFN) and pro-inflammatory cytokines that are crucial for antiviral responses and inflammation (28–30). Beyond its role in host defense, TLR7 signaling regulates various physiological processes, including B cell development and plasma cell differentiation (31). Enhanced TLR7 signaling can drive abnormal B cell survival and increase follicular and extrafollicular helper T cells (32). Aberrant TLR7 activation is associated with autoimmune disease, such as systemic lupus erythematosus (33–35). TLR7 is also functional in the neurons, where it regulates genes involved in neuronal and glial development, as well as neural activity (36, 37). As an RNA sensor, TLR7 recognizes both microbial and endogenous RNAs. Li *et al*. demonstrated that the expression of *Tlr7* is highly upregulated in embryonic HSC-enriched populations (13), raising the question of whether TLR7-mediated tonic inflammatory signaling also plays a role in embryonic hematopoiesis.

MicroRNAs (miRNAs) are a class of endogenous non-coding RNAs typically with a length of 22 nucleotides. They play crucial roles in regulating the expression of genes involved in cell proliferation, differentiation, embryonic development and immune responses (38–40). The classical function of miRNAs is to regulate gene expression post-transcriptionally by targeting mRNAs for degradation or translation repression (39, 41). Beyond their intracellular roles, microRNAs are also present in extracellular spaces, including blood plasma (42–45) and other body fluids (46). Extracellular miRNAs can act as mediators of intracellular communication in paracrine and endocrine signaling (47, 48). Notably, specific miRNAs can act as ligands for TLRs, and their binding leads to the activation of TLR signaling and the induction of immune responses (38, 49). For example, miR-21 and miR-29a can target human TLR8 (50), while specific uridine-rich extracellular miRNAs like miR-146a and miR-20 can activate TLR7 signaling through their uridine-rich motifs, triggering inflammatory cytokine production *in vitro* and *in vivo* (50–54). Mice deficient in either miR-146a or TLR7 exhibit reduced inflammation, indicating the critical role of this pathway in regulating inflammatory signaling (53). miR-146a is also highly enriched in hematopoietic cells derived from mouse embryonic stem cells (55), raising and intriguing question of whether extracellular miRNA-mediated activation of TLR7 plays a critical role in the embryonic hematopoiesis.

In this study, we elucidated the role and mechanism of TLR7 signaling in the emergence of embryonic HSCs. We identify TLR7 as an essential regulator of embryonic HSC development in zebrafish and mice. Our data showed that the endogenous microRNA miR-146a activates TLR7 signaling to drive inflammatory cytokine production via interferon regulatory factor 5 (IRF5), ultimately activating Notch signaling to promote HSC emergence. Furthermore, we demonstrate that macrophages and endothelial cells contribute to TLR7-mediated production of inflammatory cytokines and promote HSC emergence through a non-cell-autonomous mechanism. These findings establish an important role for miRNA-146a and TLR7 in activating tonic inflammatory signaling that is essential for embryonic HSC development. This study further suggests the translational potential of TLR7 agonists and miR-146a for advancing stem-cell-based therapeutics.

## Results

### TLR7 agonist promotes, and TLR7 antagonist suppresses embryonic HSC development in zebrafish

Inflammatory and innate immune signaling pathways have been shown to regulate embryonic HSC development (12, 13, 18, 20). Given the established role of TLR7 in initiating inflammation and immune response, and its upregulation in embryonic HSC-enriched populations (13), we investigated its potential function in embryonic HSC development. We used the synthetic oligoribonucleotide resiquimod (R848), a known TLR7 agonist that promotes cytokine production (56, 57), and chloroquine (CQ), which inhibits TLR7 downstream signaling by preventing transcription factor activation (58). Dechorionated zebrafish embryos were exposed to 50 µM R848 or 20 µM CQ from 11 hours post fertilization (hpf), coinciding with the onset of primitive hematopoiesis (59), until the emergence of HSCs at 30 hpf. Compared to the vehicle-treated controls, *cmyb* expression in the aorta-gonad-mesonephros (AGM) region was enhanced by R848 and reduced by CQ at 30 hpf (Figures 1A and S1A). Consistent with this, the number of *cmyb^+^*cells increased in the R848-treated Tg(*cmyb*:GFP) transgenic embryos and decreased in the CQ-treated ones (Figure 1B). The regulatory effect persisted even when drug treatment was initiated after hemogenic endothelium specification (> 24 hpf) (Figure 1C). The efficacy of the treatments was confirmed by a significant increase in *tnfa*, *il1b*, and *il6* expression following R848 exposure and a decrease after CQ treatment (Figure 1D), indicating specific activation or inhibition TLR7 signaling in the embryos.

**Figure 1.**
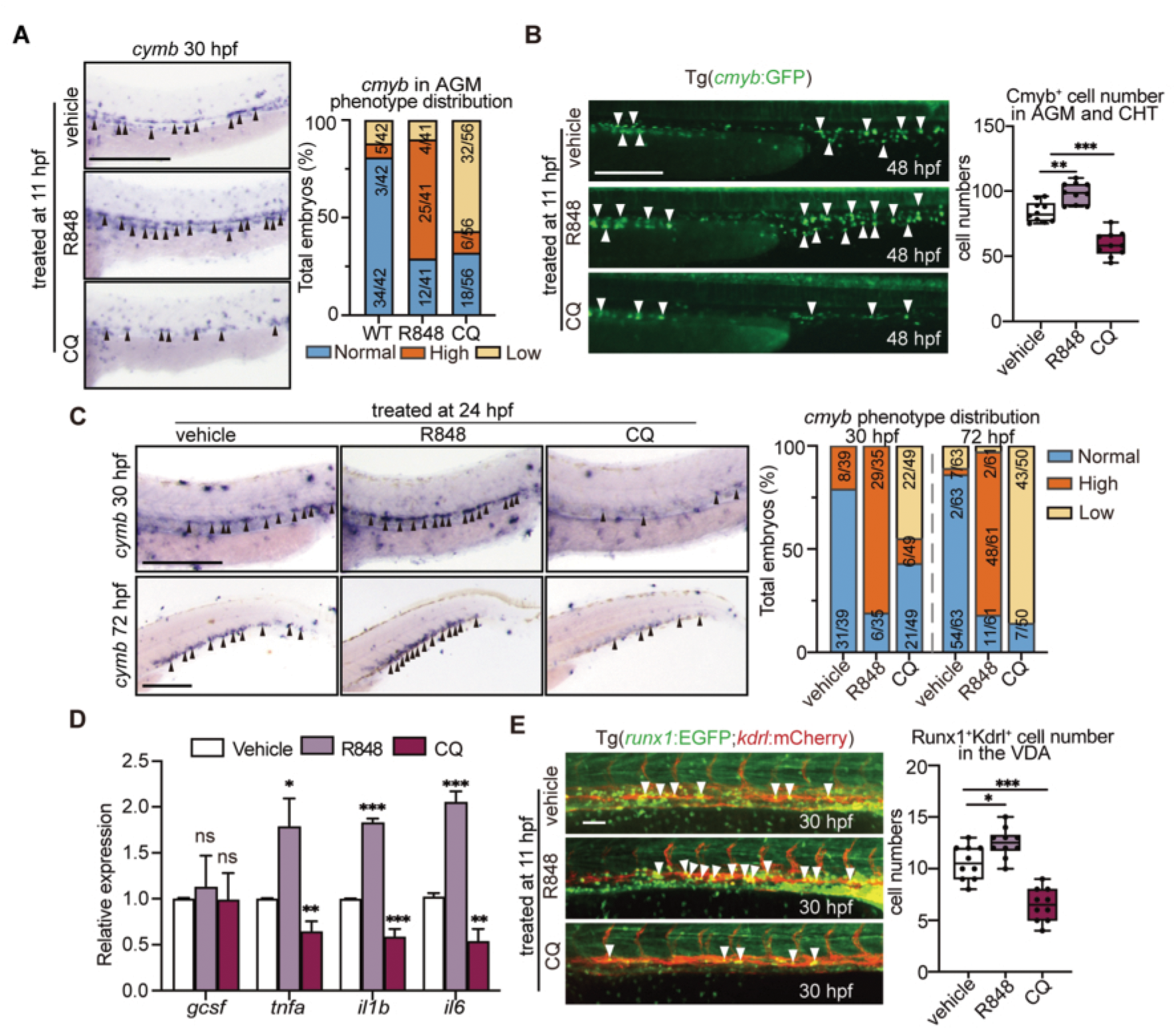
TLR7 agonist promotes, and TLR7 antagonis inhibits embryonic HSC development in zebrafish. (A) Whole-mount in situ hybridization (WISH) for *cmyb* in the aorta-gonad-mesonephros (AGM) region at 30 hours post-fertilization (hpf) (black arrowheads). Embryos were treated with vehicle control, 50 μM TLR7 agonist R848, or 20 μM antagonist Chloroquine (CQ) from 11 to 30 hpf. Its qualitative phenotypic distribution plot of embryos scored for low, high and normal *cmyb* expression. Scale bar, 200 μm. (B) Fluorescence images of hematopoietic stem and progenitor cells (HSPCs) in the AGM and caudal hematopoietic tissue (CHT) region of Tg(*cmyb*:GFP) embryos at 48 hpf (white arrowheads) following vehicle control, R848, or CQ treatment. The number of Cmyb*^+^* cells was quantified (n = 10 embryos; mean ± SD; one-way ANOVA: \*\**p* < 0.01, \*\*\**p* < 0.001). Scale bar, 200 μm. (C) WISH for *cmyb* in the AGM region at 30 hpf and in the CHT at 72 hpf (black arrowheads). Embryos were treated with vehicle control, the TLR7 agonist R848 (50 μM), or the antagonist Chloroquine (20 μM) from 24 to 30 or 72 hpf. Its qualitative phenotypic distribution plot of embryos scored for low, high and normal *cmyb* expression. Scale bar, 200 μm. (D) qPCR analysis of the inflammatory cytokine genes expression (*gcsf*, *tnfα*, *il1b,* and *il6*) in in the AGM region at 30 hpf. Embryos were treated with vehicle control, 50 μM TLR7 agonist R848, or 20 μM antagonist CQ from 11 to 30 hpf. (n = 3 biological replicates; mean ± SD; unpaired Student’s *t*-test: \**p* < 0.05, \*\**p* < 0.01, \*\*\**p* < 0.001). (E) Confocal images of hematopoietic stem cells (HSCs) in the ventral wall of the dorsal aorta (VDA) of Tg(*runx1*:EGFP;*kdrl*:mCherry) embryos at 30 hpf after treatment with vehicle control, 50 μM TLR7 agonist R848, or 20 μM antagonist CQ from 11 hpf. White arrowheads indicate Runx1^+^Kdrl^+^ cells. The number of Runx1^+^Kdrl^+^ cells was quantified (n = 10 embryos; mean ± SD; unpaired Student’s *t*-test, \**p* < 0.05, \*\*\**p* < 0.001). Scale bar, 50 μm.

To further assess the effects of TLR7 signaling modulation on HSC development, we examined HECs and EHT using Tg(*runx1*:EGFP;*kdrl*:mCherry) double-transgenic zebrafish embryos. In this model, Runx1^+^Kdrl^+^ cells mark the emergence of HSC from HECs via EHT (1, 60). Treatment with R848 from 11 hpf remarkably increased the number of Runx1^+^ and Runx1^+^Kdrl^+^ cells in the ventral wall of the dorsal aorta (VDA) at 30 hpf, whereas CQ treatment decreased their numbers (Figure S1B and 1E). Importantly, these treatments did not affect erythrocytes, endothelial cells and vascular integrity, as assessed by *gata1* and *kdrl* expression and blood flow (Figure S1C and D). Similarly, myeloid cells, such as neutrophil and macrophage, were unaffected by treatment with R848 or CQ, as indicated by *mpeg* and *mpx* expression at 28 hpf (Figure S1E). Together, these data show that activating TLR7 signaling with an agonist promotes HSC development, whereas inhibiting TLR7 signaling with an antagonist impedes it, suggesting an essential role for TLR7 signaling in embryonic HSC development.

### Generation and validation of a *tlr7*-deficient zebrafish line

To investigate the function of *tlr7* in hematopoiesis, we generated a *tlr7* mutant zebrafish line using the CRISPR/Cas9 system (Figure S2A). We designed a guide RNA (gRNA) targeting *tlr7* exon, resulting in a 4-nucleotide deletion and a subsequent premature nonsense mutation (Figure S2B). Mutant zebrafish embryos exhibited a marked reduction in *tlr7* mRNA levels in AGM cells compared to wild-type (WT) controls, likely due to nonsense-mediated decay. In contrast, mRNA levels for other related toll-like receptor genes, such as *tlr2*, *tlr4* and *tlr8*, and the adaptor *myd88*, were unchanged (Figures S2C). The WISH analysis confirmed that *tlr7* is expressed in the vessels of the AGM region at 30 hpf and the caudal hematopoietic tissue (CHT) at 72 hpf, and these expression signals were substantially diminished in *tlr7* mutants (Figure S2D), suggesting a potential role for *tlr7* in HSPC development. The successful generation of the *tlr7* mutant zebrafish line provides a valuable genetic tool for dissecting the role of TLR7 in embryonic hematopoiesis.

To valid the specificity of the chemical modulator R848 or CQ in response to TLR7 signaling, we exposed both WT and *tlr7* mutant embryos to 50 µM R848 or 30 µM CQ. Quantitative analysis of *cmyb* expression by WISH revealed that the hematopoietic-promoting effect of R848 and the suppressive effect of CQ were markedly abolished in *tlr7* mutants (Figure S2E). Collectively, the evidence indicates that the effect of R848 on promoting HSPC development is specifically dependent on TLR7.

### TLR7 is required for HSC formation but is dispensable for primitive and definitive progenitor development in zebrafish

To investigate the role of *tlr7* in HSC development, we first analyzed *cmyb* expression by WISH. At 30 hpf, *cmyb* expression was significantly reduced in the AGM region of *tlr7* mutants compared to WT embryos (Figure 2A). Consistent with this, the number of Runx1^+^Kdrl^+^ HSC cells in the VDA was remarkably lower in *tlr7* mutants within Tg(*runx1*:GFP;*kdrl*:mcherry) double-transgenic background at 30 hpf (Figure 2B). This reduction in *cmyb* expression became more pronounced at 48 and 72 hpf in *tlr7* mutants (Figure 2C). Similarly, Tg(*cmyb*:GFP) zebrafish exhibited significantly fewer GFP^+^ cells in the CHT region of *tlr7* mutants at 72 hpf (Figure 2D). The specificity of this phenotype was confirmed by the rescue of *cmyb* expression at 30 hpf upon injection of *in vitro*-transcribed *tlr7* mRNA into mutant embryos (Figure 2E). Furthermore, injection with translation-blocking morpholino (MO) targeting *tlr7* also resulted in a remarkable reduction in *cmyb* expression at 30 hpf, which was reversed by co-injection with *tlr7* mRNA (Figure 2F). These results demonstrate that TLR7 is essential for embryonic HSC development.

**Figure 2.**
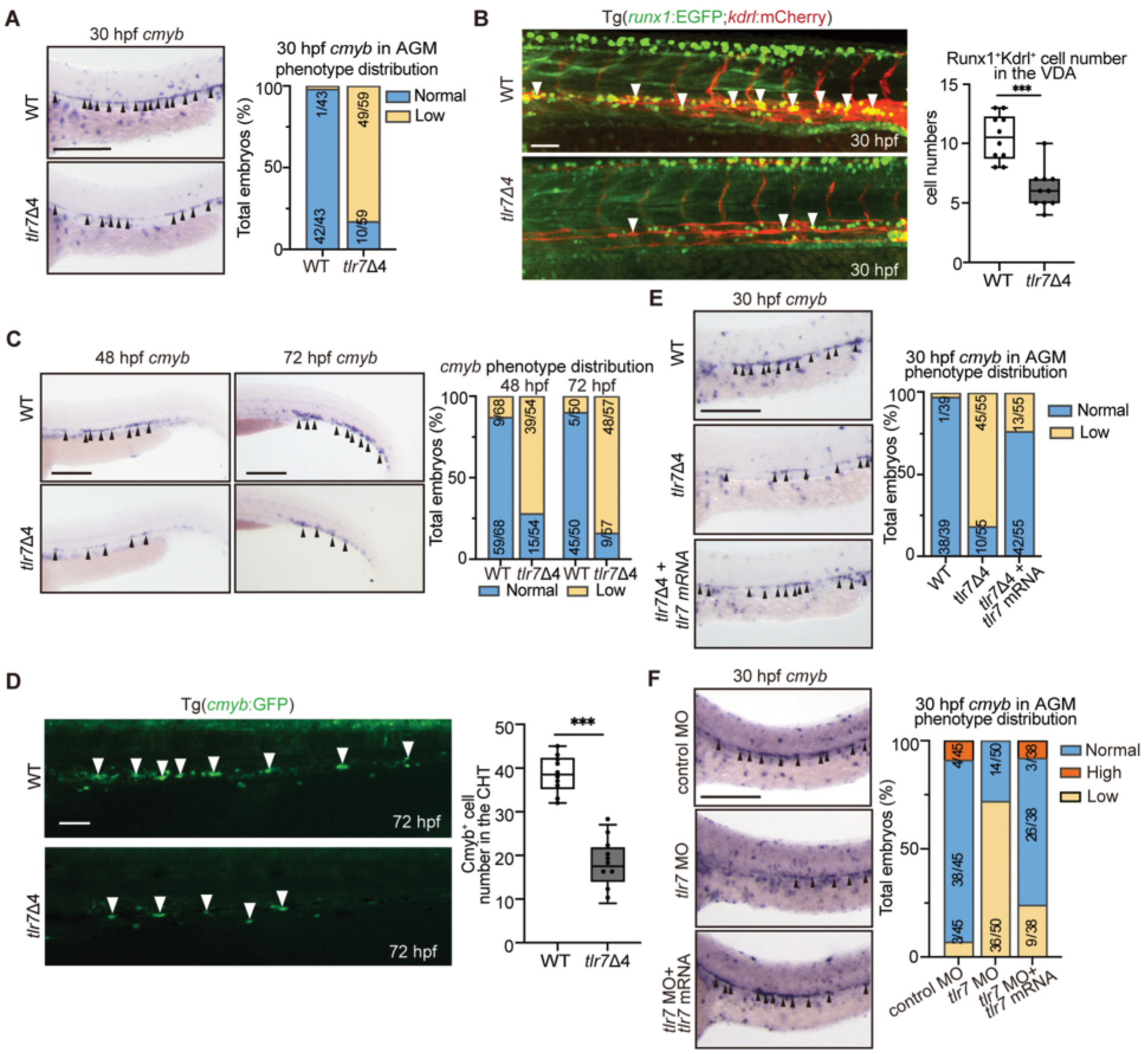
TLR7 signaling is required for HSC development. (A) Whole-mount in situ hybridization (WISH) for *cmyb* in the aorta-gonad-mesonephros (AGM) region of wild-type (WT) and *tlr7* mutants at 30 hpf (black arrowheads), and its qualitative phenotypic distribution plot of embryos scored for low or normal *cmyb* expression. Scale bar, 200 μm. (B) Confocal images of hematopoietic stem cells (HSCs) in the ventral wall of the dorsal aorta (VDA) of WT and *tlr7* mutants embryos in Tg(*runx1*:EGFP;*kdrl*:mCherry) background at 30 hpf. The number of Runx1^+^Kdrl^+^ cells was quantified (n = 10 embryos; mean ± SD; unpaired Student’s *t*-test, \*\*\**p* < 0.001). Scale bar, 50 μm. (C) WISH for *cmyb* in the AGM region at 30 hpf and in the caudal hematopoietic tissue (CHT) at 72 hpf of WT and *tlr7* mutants (black arrowheads), and its qualitative phenotypic distribution plot of embryos scored for low or normal *cmyb* expression. Scale bar, 200 μm. (D) Fluorescence images of hematopoietic stem and progenitor cells (HSPCs) in the AGM and CHT region of WT and *tlr7* mutants embryos in Tg(*cmyb*:GFP) background at 72 hpf (white arrowheads). The number of Cmyb*^+^* cells was quantified (n = 10 embryos; mean ± SD; unpaired Student’s *t*-test, \*\*\**p* < 0.001). Scale bar, 200 μm. (E) WISH for *cmyb* in the AGM region of WT and *tlr7* mutants at 30 hpf following injection of *tlr7* mRNA (black arrowheads), and its qualitative phenotypic distribution plot of embryos scored for low, normal and high cmyb expression.Scale bar, 200 μm. (F) WISH for *cmyb* in the AGM at 30 hpf following injection of *tlr7* morpholino (MO) or standard control MO, or with *tlr7* mRNA into WT embryos, and its qualitative phenotypic distribution plot of embryos scored for low, medium or high *cmyb* expression.Scale bar, 200 μm.

To determine whether *tlr7* deficiency specifically impairs EHT during HSC formation or has broader effects on hematopoiesis, we examined early progenitor and blood cell development using specific markers: *gata1* for erythroid cells, *spi1* for primitive myeloid cells, and *lcp1*, *mpx* and *mpeg* for neutrophils and macrophages. The formation of the primitive erythrocyte, neutrophils and macrophages was unaffected by *tlr7* depletion (Figures S3A-S3C). In addition, analysis of hematopoietic cells in the thymus and kidneys of adult *tlr7* mutants showed no significant differences compared to WT fishes (Figure S3D). These results suggest that TLR7 specifically regulates HSC hematopoiesis but is dispensable for primitive and definitive progenitor development.

Finally, we assessed whether general developmental defects could account for the reduced HSC in *tlr7* mutants. Vasculogenesis and blood flow in the posterior cardinal vein appeared normal in the *tlr7* mutants at 24-30 hpf (Figures S4A-S4C). TUNEL labelling at 30 hpf revealed no increase in apoptosis in the AGM or CHT region of the *tlr7* mutants (Figures S4D). These data demonstrated that the impaired HSC formation in *tlr7* mutants is not attributable to defects in vasculogenesis or cell survival, but is likely linked to a specific role of TLR7 signaling during the EHT process.

### TLR7 signaling acts through IRF5 to promote HSC development

The transcription factors IRF7 and IRF5 are well-established downstream mediators of TLR7 signaling, inducing pro-inflammatory mediators and type I interferons in innate immune cells (28–30). To determine whether TLR7 regulates HSC development through IRF5 or IRF7, we conducted loss-of-function experiments in WT zebrafish embryos using translation-blocking MO targeting *irf5* or *irf7*. WISH analysis showed that *cmyb* expression was significantly decreased in *irf5* morphants, but not in *irf7* morphants, compared to control morphants at 30 hpf (Figures 3A and S5A). Correspondingly, the number of Runx1^+^Kdrl^+^ HSCs was significantly lower in *irf5* morphants than in MO controls (Figure 3B). Consistent with a specific role for IRF5, qPCR analysis confirmed that only *irf5* expression was significantly reduced in *tlr7* mutants at 30 hpf, whereas *irf7* expression remained unchanged (Figure S5B). The hematopoietic defects in *irf5* morphants persisted at later stages, as evidenced by a significant reduction in *cmyb*^+^ HSPCs in the CHT at 48 and 72 hpf (Figures 3C and 3D).

**Figure 3.**
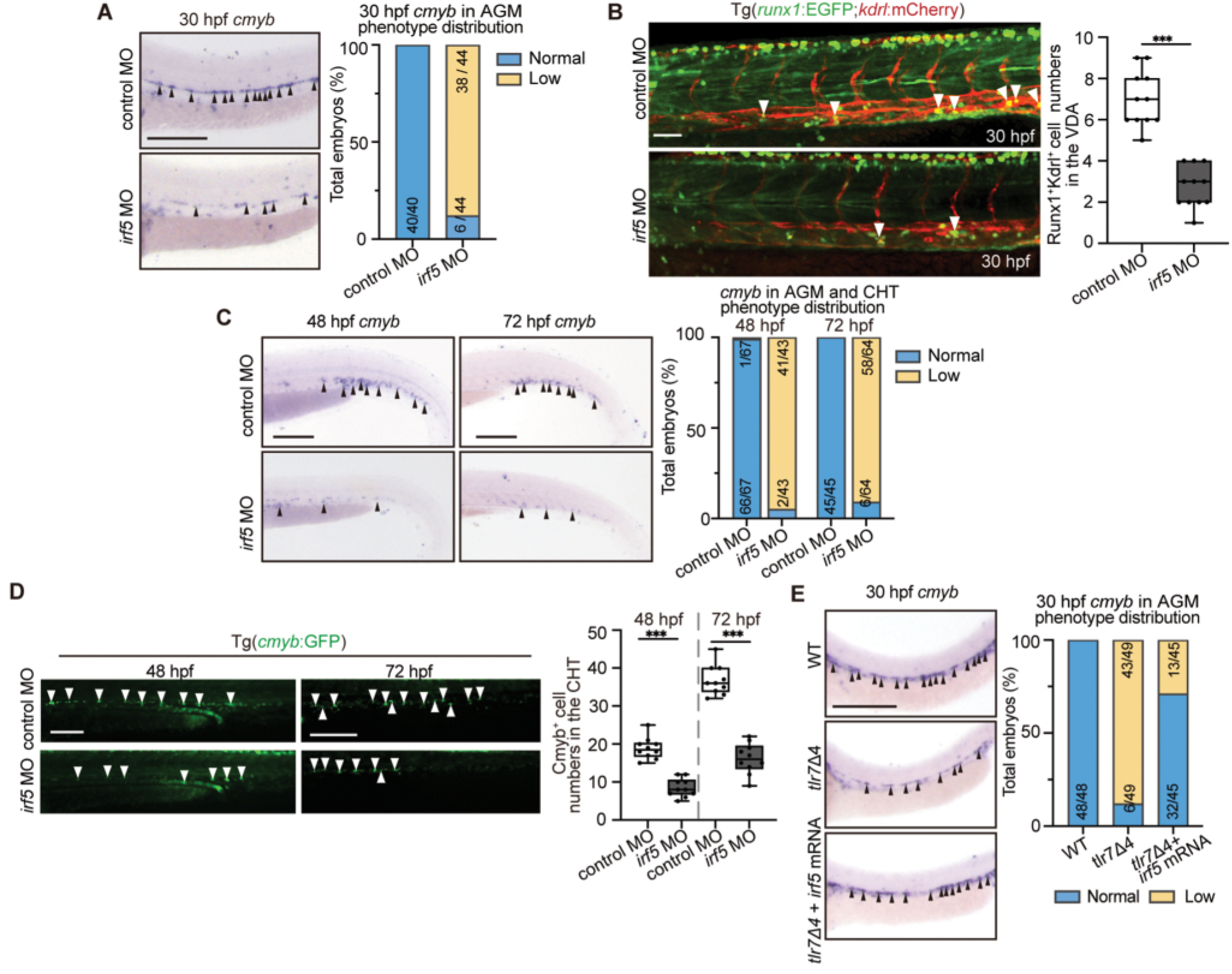
TLR7 signaling acts through IRF5 during HSC development. (A) Whole-mount in situ hybridization (WISH) for *cmyb* in the aorta-gonad-mesonephros (AGM) region of wild-type (WT) embryos at 30 hpf following injection of *irf5* translation-blocking morpholino (MO) or standard control MO, and its qualitative phenotypic distribution plot of embryos scored for low or normal *cmyb* expression. Scale bar, 200 μm. (B) Confocal images of hematopoietic stem cells (HSCs) in the ventral wall of the dorsal aorta (VDA) at 30 hpf of Tg(*runx1*:EGFP;*kdrl*:mCherry) embryos following injection of irf5 translation-blocking morpholino (MO) or standard control MO. White arrowheads indicate the Runx1^+^Kdrl^+^ cells. The number of Runx1^+^Kdrl^+^ cells was quantified (n = 11 embryos; mean ± SD; unpaired Student’s *t*-test, \*\*\**p* < 0.001). Scale bar, 50 μm. (C) WISH for *cmyb* in the caudal hematopoietic tissue (CHT) region of WT embryos at 48 and 72 hpf following injection of *irf5* translation-blocking MO or standard control MO (black arrowheads), and its qualitative phenotypic distribution plot of embryos scored for low or normal *cmyb* expression. Scale bar, 200 μm. (D) Fluorescence images of hematopoietic stem and progenitor cells (HSPCs) in the CHT region of Tg(*cmyb*:GFP) embryos at 48 and 72 hpf following injection of *irf5* translation-blocking MO or standard control MO, *irf5* morphants. White arrowheads indicates the Cmyb*^+^* cells. The number of Cmyb*^+^* cells was quantified, (n = 10 embryos; mean ± SD; unpaired Student’s *t*-test, \*\*\**p* < 0.001). Scale bar, 200 μm. (E) WISH for *cmyb* in the AGM region of WT and *tlr7* mutants embryos at 30 hpf following injection of *irf5* mRNA (black arrowheads), and its qualitative phenotypic distribution plot of embryos scored for low or normal *cmyb* expression. Scale bar, 200 μm.

To confirm these findings and exclude potential off-target effects, we used a splice-blocking MO against *irf5*. Microinjection of this MO efficiently reduced irf5 mRNA levels in the AGM region (Figure S5C) and similarly impaired *cmyb* expression at 30 hpf (Figure S5D). Notably, neither MO against *irf5* disrupted vasculogenesis, as shown by the normal expression of the arterial-specific marker *efnb2* at 28 hpf by WISH (Figure S5E). Moreover, the decreased *cmyb* expression in *tlr7* mutants and *irf5* morphants was rescued by injection with *irf5* mRNA (Figures 3E and S5E). These results indicate that IRF5 is the downstream effector of TLR7 signaling during HSC emergence.

### TLR7-IRF5 signaling promotes HSC emergence via activating Notch signaling

TLR signaling initiates an immune response via IRFs, leading to the production of inflammatory cytokines. These cytokines can act via Notch signaling to regulate RUNX1 expression and promote HSC specification (12, 20, 61–63). We hypothesized that TLR7 signaling promotes HSC emergence by upregulating inflammatory cytokine expression, thereby activating Notch signaling. To test this, we used qPCR to analyze the expression of a panel of pro-inflammatory mediators and Notch signaling components, including receptors (*notch 1a*, *notch2*, *notch3*), ligands (*jag1a*, *jag1b*, *dla*, *alb*, *dlc*, and *dld*), and targets (*hey1*, *hey2*, *her3* and *her6*), in the AGM region of *tlr7* mutants and *irf5* morphants. Compared with their respective controls, both *tlr7* mutants and *irf5* morphants showed significantly reduced mRNA levels of inflammatory cytokine genes (*tnfα*,*il1b*, and *il6*) and key Notch signaling genes (*notch1a*, *notch3*, *hey1* and *hey2*) (Figures 4A-4D). We next performed rescue experiments to determine if these downstream effectors could reverse the HSC defects in *tlr7* mutants. Injection of *tnfα* or *hey1* mRNA into *tlr7* mutant embryos efficiently restored *cmyb* expression in the AGM at 30 hpf and in the CHT at 72 hpf (Figure 4E), and also rescued the number of Runx1^+^Kdrl^+^ HSCs in the VDA at 30 hpf (Figure 4F).

**Figure 4.**
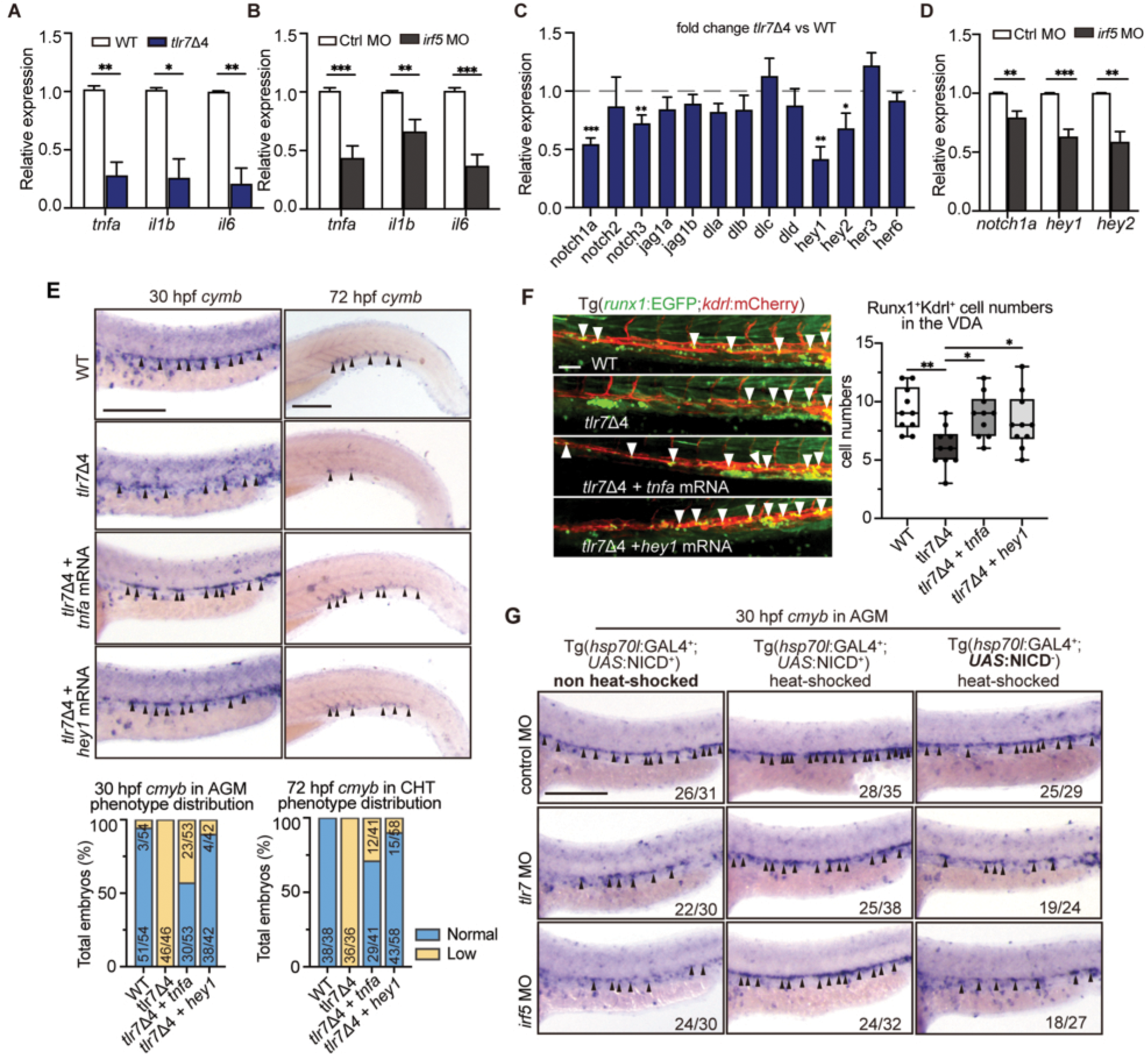
Notch signaling functions downstream of RLR7-IRF5 to regulate HSC emergence. (A) qPCR analysis of the inflammatory cytokine genes expression (*tnfα*, *il1b,* and *il6*) in the aorta-gonad-mesonephros (AGM) region of wild-type (WT) and *tlr7* mutants at 30 hpf (n = 3 biological replicates; mean ± SD; unpaired Student’s *t*-test, \**p* < 0.05, \*\**p* < 0.01). (B) qPCR analysis of the inflammatory cytokine genes expression (*tnfα*, *il1b,* and *il6*) in the AGM region at 30 hpf following injection of *irf5* translation-blocking morpholino (MO) or standard control MO (Ctrl MO) (n = 3 biological replicates, mean ± SD; Student’s *t*-test, \*\**p* < 0.01, \*\*\**p* < 0.001). (C) qPCR analysis of Notch signaling genes in the AGM region of WT and *tlr7* mutants at 30 hpf (n = 3 biological replicates, mean ± SD, Student’s *t*-test, \**p* < 0.05, \*\**p* < 0.01, \*\*\**p* < 0.001). (D) qPCR analysis of Notch signaling genes in the AGM region at 30 hpf following injection of *irf5* translation-blocking MO or standard control MO, (n = 3 biological replicates; mean ± SD; unpaired Student’s *t*-test, \*\**p* < 0.01, \*\*\**p* < 0.001). (E) WISH for *cmyb* in the AGM region of WT and *tlr7* mutants embryos at 30 hpf following injection of *tnfa or hey1* mRNA (black arrowheads), and its qualitative phenotypic distribution plot of embryos scored for low or normal *cmyb* expression. Scale bar, 200 μm. (F) Confocal images of hematopoietic stem cells (HSCs) in the ventral wall of the dorsal aorta (VDA) of WT and *tlr7* mutants embryos in the background of Tg(*runx1*: EGFP;*kdrl*:mCherry) at 30 hpf following injection of *tnfa or hey1* mRNA. White arrowheads indicate the Runx1^+^Kdrl^+^ cells. The number of Runx1^+^Kdrl^+^ cells was quantified (n = 10 embryos; mean ± SD; unpaired Student’s *t*-test, \**p*< 0.05, \*\**p* < 0.01). Scale bar, 50 μm. (G) WISH for *cmyb* in the AGM region of Tg (*hsp70l*:GAL4;*UAS*:NICD) embryos at 30 hpf following injection of *irf5* MO, *tlr7* MO, or standard control MO (black arrowheads). Scale bar, 200 μm.

To further validate Notch signaling involvement, we used the Tg (*tp1*:EGFP) reporter line, in which EGFP expression is driven by a Notch-responsive promoter (64). Confocal microscopy of Tg(*tp1*:GFP;*kdrl*:mCherry) embryo revealed markedly fewer Tp1^+^Kdrl^+^ cells in the dorsal aorta of *tlr7* and *irf5* morphants compared to control morphants at 30 hpf (Figure S6A), indicating that Notch signaling activity was attenuated upon knockdown of *tlr7* and *irf5*. Consistent with this, WISH showed reduced expression of the Notch target genes *hey1* and *hey2* in *tlr7* mutants at 30 hpf by (Figure S6B). To directly test whether Notch signaling acts downstream of TLR7-IRF5 during HSC development, we ectopically activated Notch signaling in *tlr7* and *irf5* morphants using the Tg(*hsp70l*:GAL4;*UAS*:NICD) double transgenic line, which induces expression of the Notch intracellular domain (NICD) upon heat-shock induction (65, 66). Ectopic expression of NICD effectively rescued *cmyb* expression in the AGM of both *tlr7* and *irf5* morphants at 30 hpf (Figures 4G). Collectively, these findings demonstrate that TLR7-IRF5 signaling regulates HSC emergence by activating the Notch signaling pathway.

### microRNA-146a functions as an endogenous regulator of HSC emergence via the TLR7-IRF5-Notch signaling

Since TLR7 signaling can be activated by both microbial and endogenous RNA ligands, and developing embryos are generally sterile, we hypothesized that endogenous RNAs activate TLR7 signaling during normal development. Given that miR-146a and miR-20a have been reported to activate TLR7 signaling and trigger cytokine release (50–53), and miR-146a is highly enriched in hematopoietic cells derived from mouse embryonic stem cells (55), we investigated whether TLR7 regulate embryonic HSC production by sensing these micoRNAs. Injection of miR-146a mimics, but not miR-20a mimics, significantly enhanced *cmyb* expression in the AGM at 30 hpf, compared to embryos injected with negative control microRNA mimics (NC mimics) (Figures 5A and S7A). This enhancement was absent in *tlr7* mutants but was restored upon co-injection with *tlr7* mRNA (Figure 5A). Consistent with this, the number of Runx1^+^Kdrl^+^ HSCs was significantly increased in WT embryos injected with miR-146a mimics, but not in *tlr7* mutants (Figure 5B). Conversely, injection of WT embryos with a miR-146a-specific antagomir significantly reduced *cmyb* expression levels and the number of Runx1^+^Kdrl^+^ HSCs (Figures 5C and 5D), phenocopying the HSC defects observed in *tlr7* mutants. Injection of miR-146a antagomir into *tlr7* mutant embryos produced no additional effect (Figures 5D), indicating that miR-146a functions in a *tlr7*-dependent manner. To determine if miR-146a acts through the TLR5-IRF5 axis, we injected miR-146a mimics into *Irf5* morphants. This failed to enhance HSC production (Figure 5E), suggesting that miR-146a requires IRF5 to promote HSC emergence.

**Figure 5.**
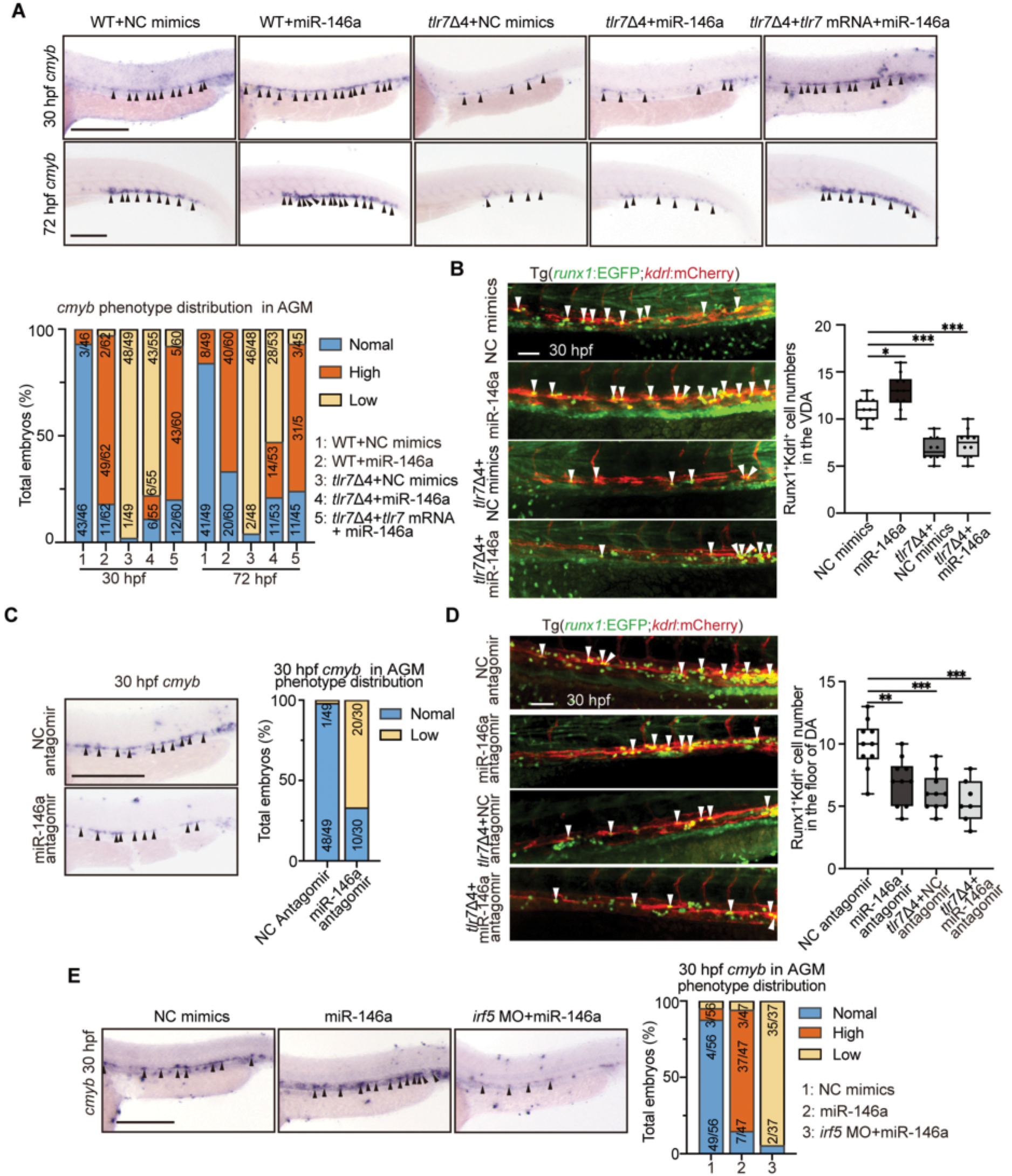
microRNA-146a functions as an endogenous regulator of TLR7-IRF5-NOTCH signaling to regulate HSPC emergence. (A) Whole-mount in situ hybridization (WISH) for *cmyb* in the aorta-gonad-mesonephros (AGM) at 30 hpf and in the caudal hematopoietic tissue (CHT) at 72hpf of wild-type (WT) and *tlr7* mutant embryos. Embryos were injected with a negative control (NC) mimics, miR-146a mimics, or miR-146a mimics co-injected with *tlr7* mRNA (black arrowheads), and its qualitative phenotypic distribution plot of embryos scored for normal, low or high *cmyb* expression. Scale bar, 200 μm. (B) Confocal images of hematopoietic stem cells (HSCs) in the ventral wall of the dorsal aorta (VDA) of WT and *tlr7* mutant embryos in Tg(*runx1*:EGFP; *kdrl*:mCherry) background at 30 hpf following injection of NC or *miR-146a* mimics. White arrowheads indicates the Runx1^+^Kdrl^+^ cells. The number of Runx1^+^Kdrl^+^ cells was quantified (n=10 embryos; mean ± SD; unpaired Student’s *t*-test, \**p* < 0.05, \*\*\**p* < 0.001). Scale bar, 50 μm. (C) WISH for *cmyb* in AGM of WT embryos at 30 hpf following injection of NC or miR-146a antagomir (black arrowheads), and its qualitative phenotypic distribution plot of embryos scored for normal, low or high *cmyb* expression. Scale bar, 200 μm. (D) Confocal images of HSCs) in the VDA of WT and *tlr7* mutant embryos in Tg(runx1:EGFP; kdrl:mCherry) background at 30 hpf following injection of NC or miR-146a antagomir. White arrowheads indicate the Runx1^+^Kdrl^+^ cells. The number of Runx1^+^Kdrl^+^ cells was quantified (n=10 embryos; mean ± SD; unpaired Student’s *t*-test, \*\**p* < 0.01, \*\*\**p* < 0.001). Scale bar, 50 μm. (E) WISH for *cmyb* in AGM of WT embryos at 30 hpf following injection of NC mimics, miR-146a mimics, or *irf5* MO with miR-146a mimics (black arrowheads), and its qualitative phenotypic distribution plot of embryos scored for normal, low or high *cmyb* expression. Scale bar, 200 μm.

Given the decreased expression of inflammatory cytokines and Notch target genes in the *tlr7* mutants and *irf5* morphants (Figures 4A-4D), we investigated the impact of miR-146a on these pathways in embryos. qPCR analysis showed that injection of miR-146a led to significant upregulation, while the antagomir caused significant downregulation, of the key transcripts within the TLR7-IRF5 signaling (*tlr7*, *irf5*, *tnfa*, *il1b* and *il6*) and its downstream Notch signaling (*notch1a*, *hey1* and *hey2*) (Figures S7B). In addition, miR-146a injection upregulated key NF-κB subunits (*nfkb1* and *p65*) in the AGM at 30 hpf (Figure S7C). These results suggest that miR-146a acts as an endogenous factor that activates TLR7-IRF5 signaling to promote HSC emergence via the Notch signaling pathway.

### The role of TLR7 signaling in embryonic hematopoiesis is conserved in mice

To determine whether the role of TLR7 signaling is evolutionarily conserved in mammalian embryonic hematopoiesis, we analyzed the CD31^+^c-Kit^+^VEC^+^ intra-arterial hematopoietic clusters (IAHCs) in mouse embryos. Genetic deletion of *Tlr7* significantly reduced the number of IAHCs in the AGM region at embryonic day 11.5 (E11.5) (Figure 6A).

**Figure 6.**
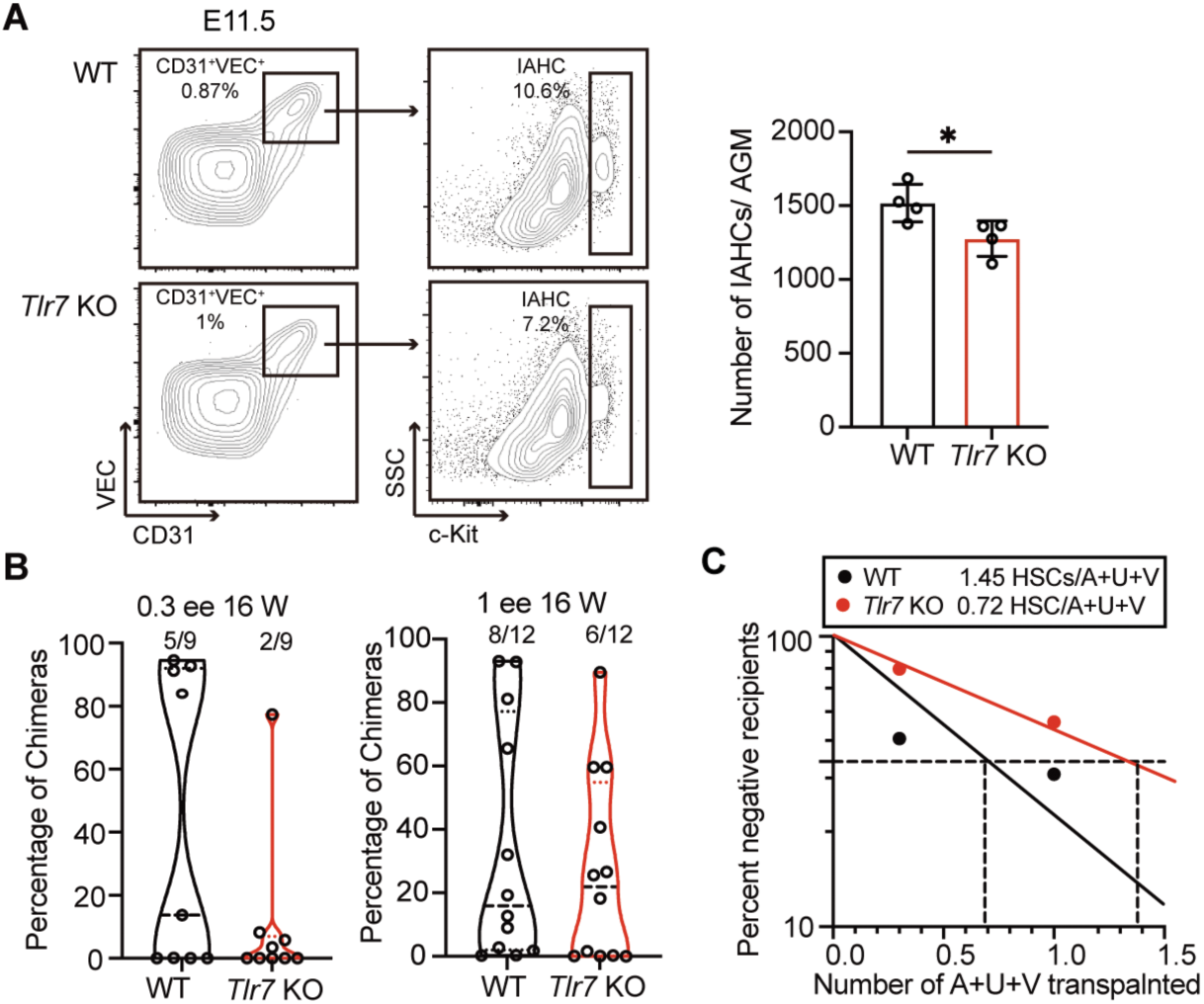
The role of TLR7 signaling in embryonic hematopoiesis is conserved in mice. (A) Representative flow cytometry plots showing the gating strategy for CD31^+^VEC^+^c-Kit^+^ intra-arterial hematopoietic clusters (IAHCs) in the aorta-gonad-mesonephros (AGM) of wilf-type (WT) and *Tlr7* knockout (KO) mouse embryos at day 11.5 (E11.5). (n = 4 biological replicates; mean ± SD; unpaired Student’s *t* test; \**p* < 0.05) (B) Chimerism analysis of peripheral blood from recipient mice after 16 weeks post-transplantation with 0.3 or 1 embryonic equivalent (ee) AGM and umbilical and vitelline arteries (A+U+V) cells from E11.5 of WT or *Tlr7* KO embryos. Data show the percentage of donor-derived cells. (C) Quantification of HSC frequency in E11.5 of WT and *Tlr7* KO mouse embryos by limiting dilution transplantation of 0.3 or 1 ee of WT or *Tlr7* KO A+U+V cells.

We next assess the number of functional HSC in the E11.5 AGM and umbilical and vitelline arteries (A+U+V) using limiting dilution transplantation assays (Figure S8A). The results showed that the engraftment potential was 2.5 times higher in recipients transplanted with 0.3 embryo equivalents (ee) WT A+U+V compared to those receiving an equal amount of *Tlr7* knockout cells (Figure 6B). There was a clear trend toward a reduced frequency of functional HSCs in *Tlr7* knockout embryos, culminating in an approximately two-fold decrease in the HSC numbers per A+U+V region compared to WT littermates (Figure 6C and S8B-8E). Furthermore, the *Tlr7*-deficient A+U+V cells contribute significantly fewer donor-derived hematopoietic cells overall in the recipient mice (Figure 6B and S8B). These results demonstrate that *Tlr7* deficiency impairs embryonic HSC development, underscoring the critical and evolutionarily conserved role of *Tlr7* in HSC emergence.

## Discussion

In this study, we identify TLR7 as a novel positive regulator of embryonic HSC development. We showed that during HSC emergence in the dorsal aorta, TLR7 stimulates the expression of inflammatory cytokine genes via IRF5 in macrophage and endothelial cells, thereby activating Notch signaling to promote HSC development through a non-cell-autonomous mechanism. Furthermore, we establish that miR-146a acts as an endogenous activator of this TLR7-IRF5 signaling to facilitate HSC emergence.

Inflammation is an evolutionarily conserved process that protects the host by activating immune and non-immune cells to eliminate pathogens and initiate tissue repair. In zebrafish and mouse models, proinflammatory cytokines such as tumor necrosis factor α (TNF-α), interferons (IFN-α and IFN-γ), and interleukins are crucial for HSPC development (12, 13, 15–19). Deficiencies in these cytokines impair HSC emergence and proliferation. A comprehensive understanding of these inflammatory signals is vital for elucidating hematological disorders and improving *ex vivo* expansion of HSCs. Toll-like receptors (TLRs) are a key family of pattern recognition receptors that mediate immune activation. Notably, transcripts for *Tlr1*, *Tlr2*, *Tlr7*, and *Tlr13* are up-regulated in HSC-enriched hematopoietic cluster cells in mouse embryos (13). Bennett *et al*. reported that loss of *Tlr2* or *Tlr4* reduces IAHCs in mouse embryos at E10.5, but increases HSCs at E11.5 (67). In this study, we found that *Tlr7* deficiency leads to a significant reduction in IAHCs at E11.5 and an approximately 50% reduction in HSCs at E11.5. The absence of a compensatory downregulation of other tlrs (e.g., *tlr2*, *tlr4*, *tlr8*) in *tlr7* mutant suggests that TLR7 functions distinctly from, and perhaps in opposition to, TLR2 and TLR4 during HSC specification in the AGM. We detected *tlr7* expression in the aorta region of zebrafish embryos, but current technical limitations precluded precise cellular localization and exact cell type. Future studies employing conditional knockout mouse models to delete *Tlr7* specifically in different cells or hematopoietic cells will be essential to determine whether TLR7 regulate HSC emergence through a cell-autonomous or non-cell-autonomous mechanism.

Canonically, TLR7 recognizes single-stranded RNA and triggers a signaling cascade via adaptor protein MyD88. Bennett *et al*. showed that loss of MyD88 decreases IAHCs but increases HSCs in the AGM region of mouse embryos (13). In our study, we found that deletion of Tlr7 severely reduces embryonic HSC numbers without affecting *myd88* expression. This indicates that TLR7 regulates HSC development through a MyD88-independent mechanism, likely involving an alternative adaptor. Recent studies have shown that SLC15A4 recruits a novel adaptor, TASL, to endolysosomes upon TLR7 activation, subsequently triggering IRF5-mediated inflammatory responses (68, 69). Whether TLR7 regulates embryonic HSC development through SLC15A4 and TASL warrants further investigation.

Following their emergence in the AGM, HSCs migrate to the fetal liver to mature and expand before seeding the bone marrow to maintain adult hematopoiesis. Although loss of TLR7 leads to reduced embryonic HSCs, *Tlr7* KO mice are viable without major overt developmental defects. This phenotype is consistent with the nature of inflammatory signaling networks, which are often complex and redundant. It is plausible that the loss of TLR7 is compensated for by other mechanisms at later developmental stages. A precedent for such stage-specific regulation exists: while embryos deficient for IFN-γ or its receptor exhibit a 2∼4 fold reduction in HSCs (13), adult HSCs from these same IFN-γ-deficient mice show enhanced engraftment in competitive transplantations (70). This highlights that distinct factors regulate HSCs from their embryonic emergence to their maintenance in the adult bone marrow (3). Our study focused on the initial emergence of HSCs from HEs and established TLR7 as essential for this process. Whether TLR7 also regulates subsequent HSC function, such as expansion, self-renewal, and differentiation, remains an open question.

MicoRNAs are crucial post-transcriptional regulators of embryonic development, including hematopoiesis. For example, miR-223 negatively regulates retinoic acid signaling, thereby repressing the generation of murine HECs and HSPCs (71). In contrast, endothelial miR-128 regulates nascent HSPC heterogeneity by promoting Wnt and Notch signaling in the AGM through post-transcriptional repression of the Wnt inhibitor *csnk1a1* and the Notch ligand *jag1b* (72). Dysregulation of miRNAs can have severe consequences, as seen in USB1 mutants, which contribute to hematopoietic failure (73). In addition, extracellular miRNAs can be sensed by TLRs to regulate various physiological and pathological processes. For instance, extracellular miR-146a promotes sepsis-induced brain inflammation via TLR7 (53, 74). A study utilizing an in *vitro* differentiation system of mouse embryonic stem cells found that miR-146a was highly enriched in CD41^+^ and/or CD45^+^ cells (55). However, the function of miR-146a in embryonic hematopoiesis was previously unknown. In this study, we showed that miR-146a mimics enhance, and miR-146a antagomirs suppress HSC development in WT zebrafish but not in *tlr7* mutants, positioning it as an upstream activator of the pathway. Future studies are warranted to examine miR-146a in the embryonic hematopoiesis by knocking out the miR-146a genes.

It is important to note that miR-146a is also a well-characterized negative feedback regulator of inflammation in adult systems, where it represses *Traf6* expression to attenuate pro-inflammatory cytokine response and maintain normal myeloproliferation in HSC quiescence in the bone marrow niche (75, 76). Although miR-146a can inhibit NF-κB in specific contexts (77, 78), we observed no such inhibition in our model; instead, we observed a modest elevation in *nfkb1* and *p65* expression. This underscores the complexity and context-dependency of miR-146a’s downstream targets. Our research identified a novel, pro-inflammatory role for miR-146a in activating tonic inflammation to promote embryonic hematopoiesis in zebrafish. Whether miR-146a also regulates mammalian embryonic HSC development warrants future investigation. In this study, we also showed that treatment of embryos with miR-146a or R848 can activate TLR7-mediated inflammatory signaling to promote HSC development in zebrafish embryos *in vivo*. Future studies will be important to investigate whether miR-146a or R848 treatment can enhance mammalian embryonic HSC development ex vivo or improve the efficiency of *in vitro* HSC differentiation from embryonic stem cells or pluripotent stem cells.

In summary, our study elucidates a critical role for TLR7-mediated inflammatory signaling in the emergence of HSC. Significantly, we identified miR-146a as an endogenous ligand that activates inflammatory signaling in healthy embryonic tissues. This finding provides new insights into the source of endogenous ligands and cytokines that contribute to tonic inflammatory signaling during normal hematopoiesis.

## Materials and Methods

### Zebrafish and mouse strains

Zebrafish strains used in this study included AB wild-type, Tg(*cmyb*:GFP), Tg(*runx1*:EGFP;*kdrl*:mCherry), Tg(*tp1*:EGFP), Tg(*hsp70l*:GAL4;*UAS*:NICD), Tg(*mpeg1*:GFP), and *tlr7* mutants. They were raised and maintained under standard conditions at 28.5°C in a recirculating aquaculture system. All zebrafish procedures were performed in accordance with the guidelines of the Laboratory Animal Center of Zhejiang University (Protocol No.: ZJU20220375).

*Tlr7* knockout mice (C57BL/6Smoc-Tlr7^em1Smoc^) on a C57BL/6 background were purchased from Shanghai Model Organisms Center, Inc. (Shanghai, NM-KO-190166). Mice were housed under standard conditions at 25°C and 50% humidity, with a 12-hour light/dark cycle. They were provided with sufficient water and food (79). All mouse procedures were approved by the Institutional Animal Care and Use Committee of the Laboratory Animal Center, Zhejiang University (Protocol No.: ZJU20230197).

### Generation of zebrafish knockout lines using CRISPR/Cas9

The zebrafish *tlr7* mutant line was generated in the wild-type AB strain background using CRISPR/Cas9 technology. A single-guide RNA (sgRNA) targeting the sequence (5’-GACTGATATCCAGAACGACCAGG-3’) within exon 1 of the *tlr7* gene was designed using the ChopChop web tool (https://chopchop.cbu.uib.no/). The gRNA template was amplified from the scaffold vector and subsequently transcribed using the T7 High Yield RNA Transcription Kit (Vazyme, TR101-01). A mixture of 4 pmol SpyCas9 protein (NEB, M0646) and 100 pg *tlr7*-targeting gRNA was co-injected into the yolk of wild-type embryos at the one-cell stage. The injected F0 founder embryos were raised to adulthood and outcrossed with wild-type zebrafish to generate F1 progeny. F1 progeny were screened for indel mutations at three months post-fertilization by PCR amplification and Sanger’s sequencing of genomic DNA from the tail fin clips. Heterozygous F1 carriers were identified and intercrossed to establish a stable homozygous F2 mutant line (80).

### Morpholino, microRNA mimic, antagomir, and mRNA injection

Translation-blocking morpholino oligonucleotides (MOs) targeting *tlr7*, *irf5*, *irf7* and *spi1b*, as well as a splice-blocking MO (sbMO) for *irf5*, were designed and purchased from Gene Tools. MO sequences were listed in Table S1. MOs were dissolved with diethyl pyrocarbonate (DEPC)-treated water to generate 1 mM as stock solutions. For microinjection, working concentrations of MOs were prepared in a solution containing phenol red as an injection tracer. Embryos were injected at the one-cell stage with the following concentrations: *tlr7* MO (0.75 mM), *irf5* MO (0.2 mM), *irf7* MO (1 mM), *spi1b* MO (0.5 mM), *irf5* sbMO (1 mM). A standard control MO was used in all experiments.

The microRNA mimic and antagomir were designed and synthesized by GenePharma. The sequences of the microRNA mimics and antagomir were listed in Table S2. Lyophilized nucleotides were reconstituted in DEPC-treated water to make a stock solution. Working solutions (miR-20a, 10 µM; miR-146a mimics, 20 µM; miR-146a antagomir, 10 µM) were prepared in phenol red and microinjected into the one-cell stage of zebrafish embryos. Injected embryos were collected at 30 hours and 72 hours post-injection. Negative control microRNA mimic and negative control microRNA antagonist were used parallelly.

The opening reading frame of *tlr7*, *irf5*, *irf7* were amplified and cloned into pCS2+ plasmid. Capped messenger RNA (mRNA) were transcribed in *vitro* using SP6 mMESSAGE mMACHINE^TM^ mRNA transcription synthesis kit (Invitrogen; AM1344) and purified by lithium chloride precipitation. For microinjection, mRNA was diluted to 100 ng/μL, loaded into a microinjection needle, and approximately 2 nL was delivered into one-cell-stage embryos. Following injection, embryos are transferred to a petri dish containing Danio buffer and incubated at 28.5°C.

### Whole-mount in situ hybridization

Embryos were fixed in 4% paraformaldehyde (PFA) in phosphate-buffered saline (PBS) overnight at 4°C. Digoxigenin (DIG)-labelled RNA probes for *cymb, kdrl, spi1b, gata1, hey1, hey2, mpx, mpeg1.1 and tlr7* were transcribed *in vitro* using T7 polymerase and labelled with DIG RNA labelling Mix (Roche; 11277073910). Whole-mount in situ hybridization (WISH) was performed as described previously. Following hybridization with probe, embryos were incubated with anti-digoxigenin Fab fragment antibody (Roche, 59318423), followed by incubation with 5-bromo-4-Chloro-3-indolyl-phosphate nitro blue tetrazolium (Roche, 65752120) to develop color. Embryos exhibiting specific staining were imaged, and gene expression levels were quantified by categorizing signal intensity or number of positive-staining cells as high, medium or low relative to the median expression of sibling controls (81). Only embryos that met standard morphological criteria for normal development were selected for quantitative analysis.

### Confocal microscopy

Transgenic embryos, including Tg (*cmyb*:GFP), Tg(*runx1*:EGFP;*kdrl*:mCherry) double-transgenic lines, were anesthetized and mounted in 0.8% low-melting agarose or methylcellulose in glass-bottom dishes (Cellvis, D35-20-0-N). Confocal z-stacks of the head, dorsal aorta (DA) or caudal hematopoietic tissue (CHT) region were photographed using confocal laser scanning microscop-y (Olympus CSU-W1) with a 20x objective. Z-stacks were collected at a step size of 2 µm. The image analysis was performed using Olympus confocal software, OlyVIA.

### Chemical treatment

Zebrafish embryos were treated with Resiquimod (R848) and Chloroquine (Cq) from the 11-somite stage until 30 hpf,48 hpf and 72 hpf. Wild-type embryos were incubated in 50 µM R848 or 30 µM Cq, or 0.1% DMSO (vehicle control) in Danio buffer.

### TUNEL assay

The TUNEL assay was performed using the fluorescein-based in Tunel Brightred Apoptosis Detection Kit (Vazyme, A113) in accordance with the manufacturer’s instructions. Briefly, manually dechorionated embryos were fixed in 4% PFA overnight at 4°C. After fixation, embryos were washed three times with PBST and dehydrated in 100% methanol at −20 °C for at least 2 hours. The embryos were then gradually rehydrated, washed again with PBST, and permeabilized with proteinase K and acetone. Following three additional washes in PBST, the permeabilized embryos were incubated with the labeling solution at room temperature for 30 min. Finally, embryos were washed 3 times with PBST and then were imaged by confocal microscopy.

### Heat-shock treatment

To induce the expression of Notch intracellular domain (NICD), zebrafish embryos carrying the Tg(*hsp70l*:Gal4;*UAS*:NICD) transgene were heat-shocked at 37 °C for 50 minutes at 18 hours post-fertilization (hpf). After heat shock, the embryos were transferred back and maintained at 28.5 °C until the specified time of collection.

### Limiting dilution transplantation of mouse embryonic cells

Eight-week-old CD45.1/CD45.2 female recipient mice were subjected to a split dose of 9 Gy X-ray irradiation, administered 24 hours apart. Each irradiated adult recipient was injected retro-orbitally 1.0 or 0.3 embryo equivalents (ee) of CD45.2^+^ cells isolated from aorta-gonad-mesonephros (AGM) and umbilical and vitelline arteries (A+U+V) cells from embryonic day 11.5 (E11.5) embryos (46-48 somite pairs), along with 5 × 10^4^ CD45.1 nucleated bone marrow cells as a carrier. Donor engraftment (CD45.2^+^) was assessed in peripheral blood and bone marrow 16 weeks post-transplantation. Hematopoietic stem cell (HSC) frequencies were determined using extreme limiting dilution analysis (ELDA) (82).

### Flow Cytometry

To measure the intra-arterial hematopoietic clusters (IAHCs), E11.5 aorta-gonad-mesonephros (AGM) regions were microdissected and dissociated with collagenase (Sigma; C0130). Cells were stained with antibodies to PerCP-Cy5.5 CD31 (145-2C1), PE VEC (eBioBV13), APC-Cy7 c-kit (2B8).

To analyze donor-derived cells post-transplantation, peripheral blood were collected after 4, 8, 12, and 16 weeks post-transplantation. Red blood cells were lysed using red blood cell lysis buffer (Sigma; R 7757), and remaining cells were stained with antibodies: PE CD3ε (145-2C1), APC CD19 (1D3), APC-Cy7 CD11b (M1/70), PerCP-Cy5.5 Gr-1 (RB6-8C5), PE-Cy7 CD45.1 (A20), FITC CD45.2 (104). For the detection of donor-derived long-term HSCs from bone marrow cells after 16 weeks post-transplantion, the femur and tibiofibular were collected and crushed to isolate bomarrow cells. Following red blood cell lysis, the cells were stained with antibodies: FITC anti-CD45.2 (104), PE-Cy7 anti-CD45.1 (A20), PerCP-Cy5.5 anti-Sca1 (D7), APC-Cy7 anti-c-kit (2B8), eFluor 450 lineage cocktail [(anti-TER119 (TER119), anti-Gr-1 (RB6-8C5), anti-B220 (RA3-6B2), anti-CD3 (17A2), and anti-Mac-1 (M1/70)], APC anti-CD48 (HM48-1), PE-Cy7 anti-CD150 (TC15-12F12.2). DAPI was used to exclude dead cells.

All antibodies were purchased from eBioscience (ThermoFisher Scientific). Cells were analyzed by flow cytometer ACEA NovoCyte (Agilent). Data were analyzed with FlowJo software.

### AGM explant culture

AGM dissected from E10.5 murine embryos (36-38 somite pairs) were cultured for 72 h at 37°C in a 5% CO_2_ atmosphere at the air-liquid interface. The AGM explants were maintained in six-well plates on a metal mesh covered with a Durapore membrane filter (Millipore, DVPP02500). The culture medium consists of Myeloid Long Term Culture Media (StemCell Technologies, M5300) supplemented with 10^−6^ M Hydrocortisone hemisuccinate (StemCell Technologies, 74142), 100 ng/mL SCF, 100 ng/mL IL-3 and 100 ng/mL Flt3 ligand (Novoprotein, C775, CP39 and CC19). AGM explants were pulsed with 50 μM R-848 or vehicle control during the first 2 hours of culture.

### Real-time quantitative PCR (qPCR)

Total RNA was extracted from pools of zebrafish AGMs using the Easy RNA Extraction Kit (according to the manufacturer’s instructions (Easy-do Bio, DR0401050). Then, 1 µg of total RNA was reverse-transcribed into cDNA using HiScript II Q Select RT SuperMix (Vazyme, 7E0982K4). qPCR was performed using the ChamQ Universal SYBR qPCR Master Mix (Vazyme, R323-01) on a LightCycler^®^ 480 II quantitative PCR system (Roche). Gene expression levels were normalized to *ef1a* as an internal control and calculated using the 2^-ΔΔCt^ method. Each experiment included at least three biological replicates, and results were presented as mean ± standard deviation. The sequence of all primers used is listed in Table S3.

### Bioinformatics analysis

The published single-cell RNA-seq data from mouse embryos were analyzed using an online tool (https://omg.gs.washington.edu/jax/public/index.html) (83).

## Statistical analysis

All statistical analyses were performed using GraphPad Prism (version 9). Data are presented as the mean ± standard deviation (SD). Sample sizes for each experiment are indicated in the figure legends. For comparisons between two groups, a unpaired Student’s *t*-test was used. For comparisons among more than two groups, one-way ANOVA was applied. Statistical significance was defined as follows: * *p* < 0.05, ** *p* < 0.01, *** *p* < 0.001; n.s., not significant.

## Acknowledgments

We thank Yingniang Li, Yingying Huang, and Jiajia Wang from the Core Facilities, Zhejiang University School of Medicine for their technical support. We thank Yulan Jin, Xueqiu Chen and Weidong Zeng from the Experimental Teaching Center, Zhejiang University College of Animal Sciences, for using the qPCR instrument (LightCycler^®^ 480 II, Roche). We thank Dr. Ying Shan at the Core Facility Platform of the College of Animal Sciences, Zhejiang University, for providing technical assistance on confocal imaging. We thank Dr. Pengxu Qian and Dr. Ruxiu Tie for their support.

This work was supported by the National Program on Key Research Project of China (2019YFE0103900), the National Natural Science Foundation of China (31872837, 32102620), and Hangzhou Chengxi Sci-tech innovation Corridor Management Committee.

